# Multiscale topological analysis of kinetoplast DNA via high-resolution AFM

**DOI:** 10.1101/2024.09.06.611637

**Authors:** Bradley Diggines, Sylvia Whittle, Indresh Yadav, Elizabeth P. Holmes, Daniel E. Rollins, Thomas E. Catley, Patrick S. Doyle, Alice L.B. Pyne

## Abstract

Kinetoplast DNA is a complex nanoscale network, naturally assembled from thousands of interconnected DNA circles within the mitochondrion of certain parasites. Despite the relevance of this molecule to parasitology and the recent discovery of tuneable mechanics, its topology remains highly contested. Here we present a multiscale analysis into the structure of kDNA using a combination of high-resolution atomic force microscopy and custom-designed image analysis protocols. By capturing a notably large set of high-resolution images, we are able to look beyond individual kDNA variations and quantify population properties throughout several length scales. Within the sample, geometric fluctuations of area and mean curvature are observed, corresponding with previous in-vitro measurements. These translate to localised variations in density, with a sample-wide decrease in DNA density from the outer rim of the molecule to the centre and an increase in pore size. Nodes were investigated in a single molecule study, and their estimated connectivity significantly exceeded mean valence, with a high dependence on their position in the network. While node separation was approximately half the minicircle circumference, it followed a strong bimodal distribution, suggesting more complex underlying behaviour. Finally, upon selective digestion of the network, breakdown of the fibril-cap heterogeneity was observed, with molecules expanding less upon immobilisation on the mica surface. Additionally, selective digestion was seen in localised areas of the network, increasing pore size disproportionately. Overall, the combination of high-resolution AFM and single molecule image analysis provides a promising method to the continued investigation of complex nanoscale structures. These findings support the ongoing characterisation of kDNA topology to aid understanding of its biological and mechanical phenomena.

## Introduction

Accurately describing the topological states of a nanomaterial is a vital prerequisite to the engineering of properties and function. While natural biological structures can provide significant inspiration to modern nanoscience (1–4), their multi-faceted complexity often makes effective characterisation difficult. This is no more apparent than in the kinetoplast DNA (kDNA) genome. Formed within the mitochondrion of certain unicellular parasites of the *Trypanosomatidae* family (5–7), which are responsible for up to 10000 deaths per year via diseases such as African sleeping sickness (8), kDNA is a 2D catenated mesh of DNA circles interconnected like mediaeval chainmail. Within the network, there is a combination of several thousand small DNA ‘minicircles’ (2500 bp) and up to fifty larger ‘maxicircles’ (30000 bp), which pack into dense layers in-vivo, approximately 0.5 μm thick. On release from confinement in-vitro, kDNA assumes a cupped conformation with a wavy edge, resembling a jellyfish (9). However, due to the mechanical flexibility of DNA and high packing density, the DNA circles tangle and overlap each other, creating a seemingly disordered circular web.

Early work on kDNA used bulk methods to investigate topology, with Chen et al. applying digestion and gel electrophoresis to estimate the coordination of DNA circles to be 3 (10,11). This was soon followed by nanoscale imaging, which revealed the topology of kDNA to be more heterogeneous than expected. Although the first images applied electron microscopy (EM) (12), the recent use of atomic force microscopy (AFM) has been most insightful and hints towards complex, yet ordered topologies (13–15). To prepare for AFM imaging, the kDNA is adhered and dried to the surface, inducing flattening and expansion, which reduces the extent of overlap of DNA minicircles. The first AFM study by Cavalcanti et al. (13) revealed a bi-heterogeneous structure, with a high-density fibril outlining the molecule cap. Along this fibril, many dense node sites are observed, with DNA minicircles branching out in rosette patterns. Whilst heterogeneity in the morphology of individual molecules has been observed, by subjecting these molecules to external forces, consistent and predictable behaviour is seen (16). In slit-like confinement, expansion occurs consistent with the Flory model of a 2D polymer and regular circular morphologies are assumed (17). It was recently shown that, by selectively removing particular DNA circles with restriction enzymes, it is possible to achieve tunable control of relaxation behaviour (18). He et al. linked AFM images with MD simulations (15), confirming coordination distributions and theorising the formation of nodes via minicircle chain buckling upon flattening. The regularity of these behaviours appears at odds with the seemingly disordered topology of individual DNA strands at the nanoscale.

Despite the insights of these investigations, they have fundamentally been limited by the quality of AFM imaging and the lack of in-depth analysis. Whilst the appearance of kDNA molecules have displayed consistent motifs, sub-optimal immobilisation and drying conditions result in distorted molecules with desiccation patterns as a result of chain coagulation. This, and/or low-resolution imaging prevents the thorough analysis of nano-topological details which are seemingly critical to the mechanics and biological function of kDNA. Additionally, low sample sizes of 1-2 molecules limit potential identification of inter-molecule trends and outlying individual character. Here, by applying high-resolution AFM over a large sample of kDNA molecules and implementing a custom automated analysis pipeline, we present a complete study aiming to link nano-scale topology to sample-wide molecule trends. By optimising immobilisation procedures, and linking with high resolution AFM, we are able to image kDNA to an excellent standard over many molecules. Additionally, ensuring that only the highest quality images are selected in quantitative studies provides the consistency to analyse large datasets of an otherwise variable system. Through the development of custom-built image analysis tools aided by the TopoStats software (19), we are able to quantify and characterise these complex networks over a range of length-scales. Finally, we use this pipeline to investigate the effect of enzymatic digestion on the structure of the kDNA.

## Results and Discussion

### High-resolution imaging reveals multiscale topology in kDNA molecules

Optimisation of the immobilisation and drying conditions of the kDNA onto the mica surface allowed for consistent high-resolution imaging of the kDNA network (Fig. 1 and Fig. S1a-c). For larger scale images, we use increased sampling of 1600 pixels per image or ∼10 nm per pixel, which enables us to perform quantitative analysis of the entire structure. We are able to resolve individual DNA strands (3-6 nm wide) which gives the network a more spindly appearance than has been noted in previous AFM studies (13,15) (Fig. 1b-d). It is also possible to resolve the major nodes which line the fibril at the edge of the molecule (Fig. 1c and d), in addition to a series of smaller nodes which sit within the interior cap (Fig. 1b). Whilst the majority of the immobilised molecules were in a flat and open conformation, there were also instances where the fibril folded over the cap (Fig. S1(d-f)). Due to the large size of the kDNA molecule in comparison to the effective width of the individual DNA strands, high-resolution (∼3 nm per pixel) images of subsections of the molecules were acquired to allow complete analysis at all length-scales. Sampled regions of the cap (Fig. 1b), fibril (Fig. 1c) and nodes (Fig. 1d) were taken in addition to high resolution scans of the whole molecule. This allows the observation of new structural characteristics, including the overlap and intertwining of DNA strands, single molecule density distributions and texture of node sites.

**Fig. 1.**
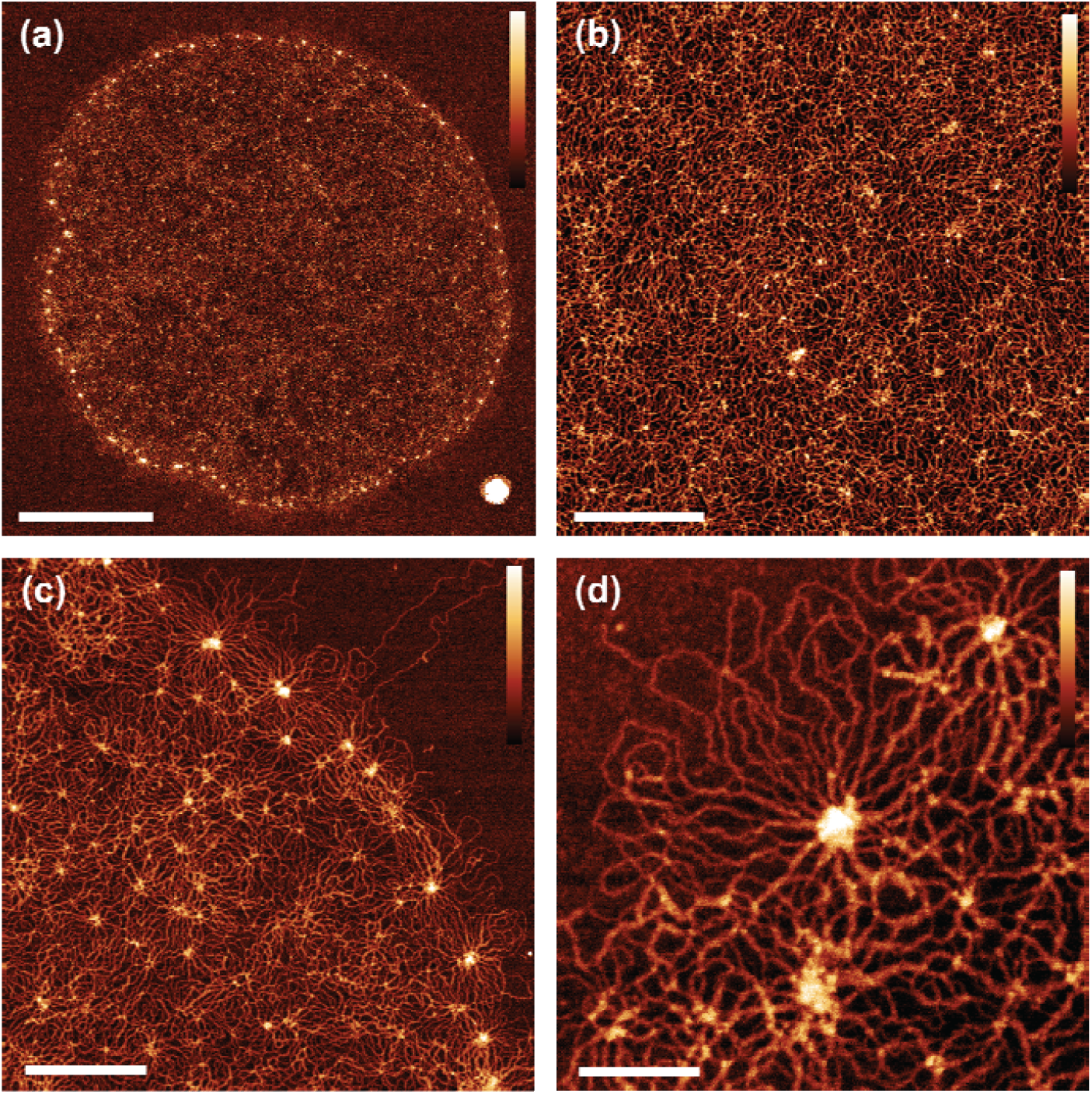
AFM images of surface-immobilised kinetoplast DNA (kDNA) captured at high resolution. (a) Whole molecule imaged with a resolution of 5 nm/px. Scale bar: 2 μm. (b) Section of interior cap with small interior nodes. Scale bar: 500 nm. (c) Fibril with a series of large nodes. Scale bar: 500 nm. (d) Single fibril node Scale bar: 150 nm. For all images Height scale: 3 nm.

### Characterisation of cap network structure

Explaining the large-scale characteristics of mechanics and geometry requires intra-molecular analysis, where a shorter length-scale is probed. Earlier studies began this by analysing fibril-cap heterogeneity and density distributions, yet these investigations were limited by sample size or quality of AFM performed (13,15). Such measurements are highly sensitive to imaging techniques, and factors such as tip size and contamination give a high likelihood for systematic error. To begin the characterisation of the network, we visualise the mean pixel height distribution for a single kDNA molecule (Fig. 2a(i)). As established in previous works, the fibril node sites around the edge of the molecule are where the greatest DNA density is located. This is further confirmed with an average of 15 molecules (Fig. 2a(ii)), which shows a peak in the mean height at the fibril edge followed by a steady decrease towards the centre of the molecule. In the negative direction - beyond the edge of the fibril - a sharp decrease is seen, corresponding to the bounds of the molecule and detection of the mica plane. While a steady gradient change is observed in bulk, individual character to the density distribution around the fibril is present with dense regions that appear to ‘snake’ inwards from the outer fibril, possibly originating from high curvature sites along the fibril.

**Fig. 2.**
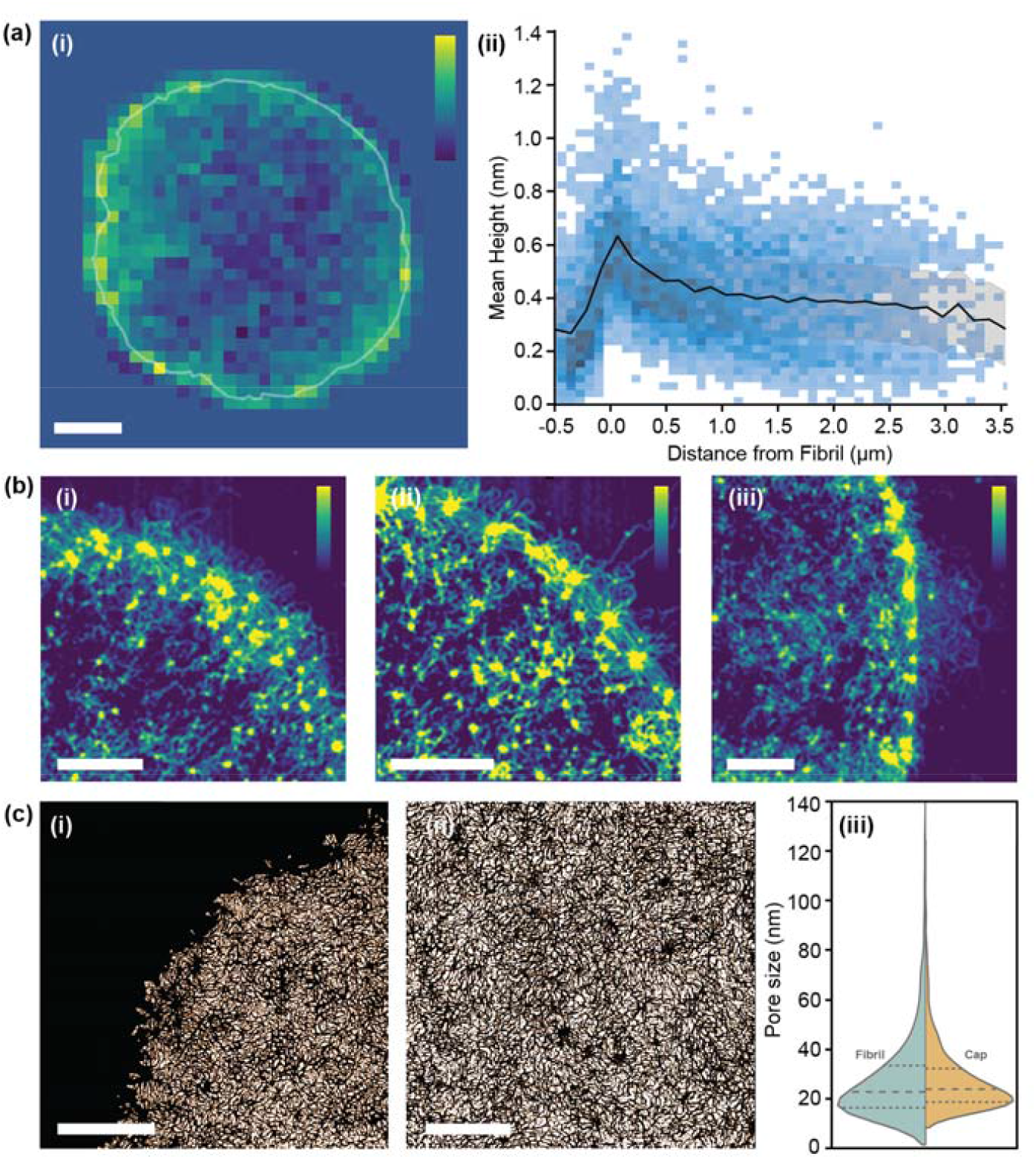
Network-wide structural characteristics of kDNA molecules. (a) **i**. Mean image height of single molecule with 300 nm sample resolution sample grid. Scale bar: 2 μm, Height scale: 2nm **ii**. Mean image height against distance from fibril for 15 molecules. (b) High contrast height images of close-ups of the outer fibril. Scale bar: 200 nm. Height scale: 1-4nm, (c) **i-ii**. Binary images highlighting pores in the network. Scale bar: 500 nm, **iii**. Pore size distribution for near-fibril and interior cap.

Such density heterogeneity is best visualised on small scale images (Fig. 2b), where a high contrast filter is applied to accentuate the most dense DNA areas. Here we see DNA pathways, dotted with a series of internal nodes (bright spots), trailing from the outer fibril towards the centre of the molecule. Interestingly, these internal trails often appear to correlate with fibril or external features, such as the DNA ‘nest’ on Fig. 2b(iii). This potentially aligns with theories for the replication of kDNA, which suggest that the greatest topoisomerase activity occurs along the outer boundaries (20).

A character of the network closely linked to the mechanical properties of 2D sheets is pore size. Instead of taking a segmentation approach to calculating this (15), we take an absolute measure of pore gaps between DNA strands (Fig. 2c). By using this method, we ignore dense regions of DNA, such as nodes, which would otherwise be included in the pore measurement. Evaluating a sample of 6 images over 4 molecules, we find a mean pore size in the cap of 28.3 ± 16.4 nm. Comparatively, for the fibril, a pore size of 26.7 ± 16.2 nm was found, consistent with findings by He et al. (34.0 ± 16.6 nm) (15). While cap and fibril have similar values, a Welch’s t-test relation of 2.42 implies sufficient statistical difference (□=0.05) and results correspond with the trends in density observed - with more dense regions producing smaller pores. These measurements confirm the heterogeneous nature of the network both in the cap and at the fibril, however further analysis is required to understand how the fibril relates to the overall structure of the molecule.

### Automated quantification of molecule global morphology

At the largest length scale, morphological analysis provides an insight into the whole molecule geometry. As observed in-vitro (9), molecules continue to show variation in edge geometry after immobilisation onto the mica surface (Fig. S1(a-c)). To quantify this over a larger sample an automated image analysis pipeline using the open-source software TopoStats, was developed. In this procedure, the exterior fibril nodes are first detected, then filtered and connected to form an outline of the molecule (Fig. 3a). Flattened molecules (N = 11) were measured to have a mean Feret diameter in the range of 7.01 μm to 8.03 μm, ± 0.05 μm, with an average of 7.6 ± 0.1 μm (Fig. 3b), consistent with previous measurements by Cavalcanti (13). Given the mean in-vitro diameter of ∼5 μm (9), this demonstrates the occurrence of significant expansion upon immobilisation of up to 60%. After immobilisation, molecules have a closely distributed perimeter length of 24.6 ± 0.1 μm (Fig. 3c), suggesting that the physical resistance of the outer fibril regulates the extent of expansion. Despite this consistency, adhered molecules still assume a range of flattened conformations with individual character. This could be due to differences in the inherent topology of kDNA molecules and/or the statistical fluctuation responsible for molecular individualism even for perfectly identical macromolecules (21). The cap area has a broad distribution, measuring 43.1 ± 5.5 μm^2^ (Fig. 3c). This variance generally correlates to the characteristics of the fibril; from smooth and wide (large area) to frilled and collapsed (low area).

**Fig. 3.**
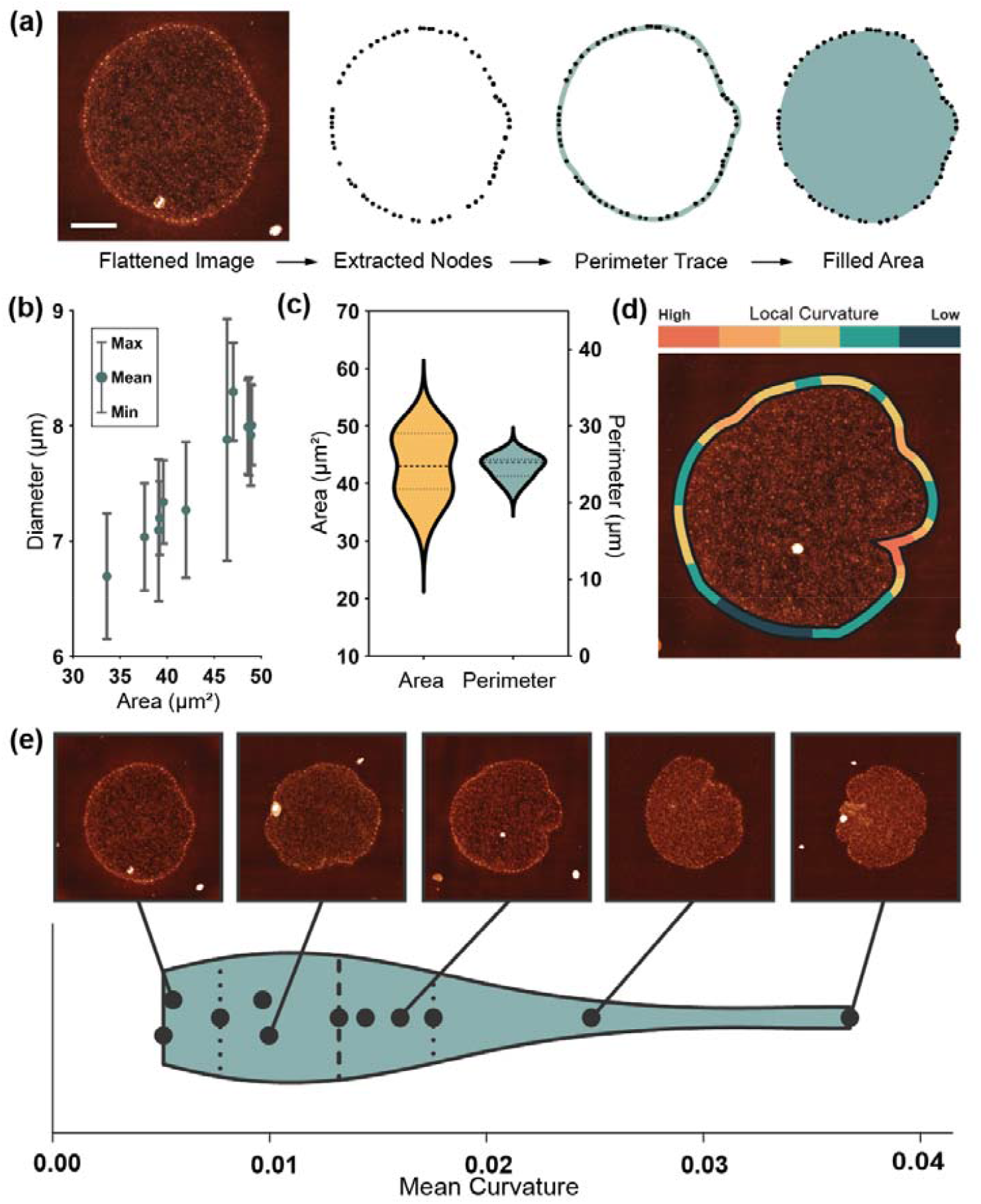
Quantifying global structure (a) Procedure to extract node positions and geometric data from flattened images. Scale bar: 2 μm (b) Molecule diameter (value bars correspond to the maximum and minimum Feret diameter of each molecule) against the measured surface area. (c) Distribution of area and perimeter for a sample of 16 molecules. (d) Graphical representation of local curvature over a single kDNA molecule (e) Violin plot showing the distribution of mean molecule curvature over 11 molecules, with corresponding images at positions along the scale.

To quantify this edge behaviour more accurately, we took local curvature measurements along the fibril length (Fig. 3d). These curvature measurements map intuitively, with such open smooth structures having a low mean curvature, and those with large inlets and frills, a high curvature. Curvature-mapped molecules can also visualise local deformation, relating features such as inlet sites to high curvature. The mean curvature over 11 molecules was 0.012, suggesting a smooth, open shape with small, localised deformations to be most common. This correlates with observations by Dolye et al. (17) during confinement, again suggesting that the immobilisation flattens, expands and smooths the network. Molecules adhered and dried on the surface for AFM measurement have a diameter of ∼7 μm, similar to that measured in confinement, but they maintain more of their anisotropic character as seen in-vitro. Additionally, we find kDNA to be immobilised on the mica surface in distinct adhesion states, a phenomenon not reported previously. Approximately half of molecules are seen lying conventionally flat on the surface, with the fibril surrounding the cap, but an equal number are observed in a stage of folding, with the fibril located either above or below the cap (Fig. S1). Since this is likely a phenomenon driven by the immobilisation kinetics, these molecules are not of interest for differing topological arrangement and are not included in further studies. However, it provides a potentially interesting insight into the mechanical properties of kDNA, for it would seem unlikely for the flattened state to be a preferred conformation. This suggests, in combination with the consistent perimeter length, that the mechanical resistance of the fibril may have a significant role in the flattening and expansion of the molecule over the surface.

### Automated determination of node connectivity

Quantification of nanoscale network configurations is inherently difficult in kDNA due to the density of overlapping strands. Even with very high-resolution AFM, the effective DNA width is 4-6 nm which is often close to that of the strand separation. This leads to problems identifying and tracing individual strands in the network. To overcome this, we first analyse a specific characteristic of the network - node sites. Located on both the cap and fibril, they present a uniquely ordered environment where strands align radially outwards in rosette-like patterns. Using an automated analysis procedure, we are able to count the number of branches around a node using a circular mask (Fig. 4a and Fig. S2). These branches often are made up of more than two adhered strands, so an area-based correction is applied to estimate the true number of connected minicircles.

**Fig. 4.**
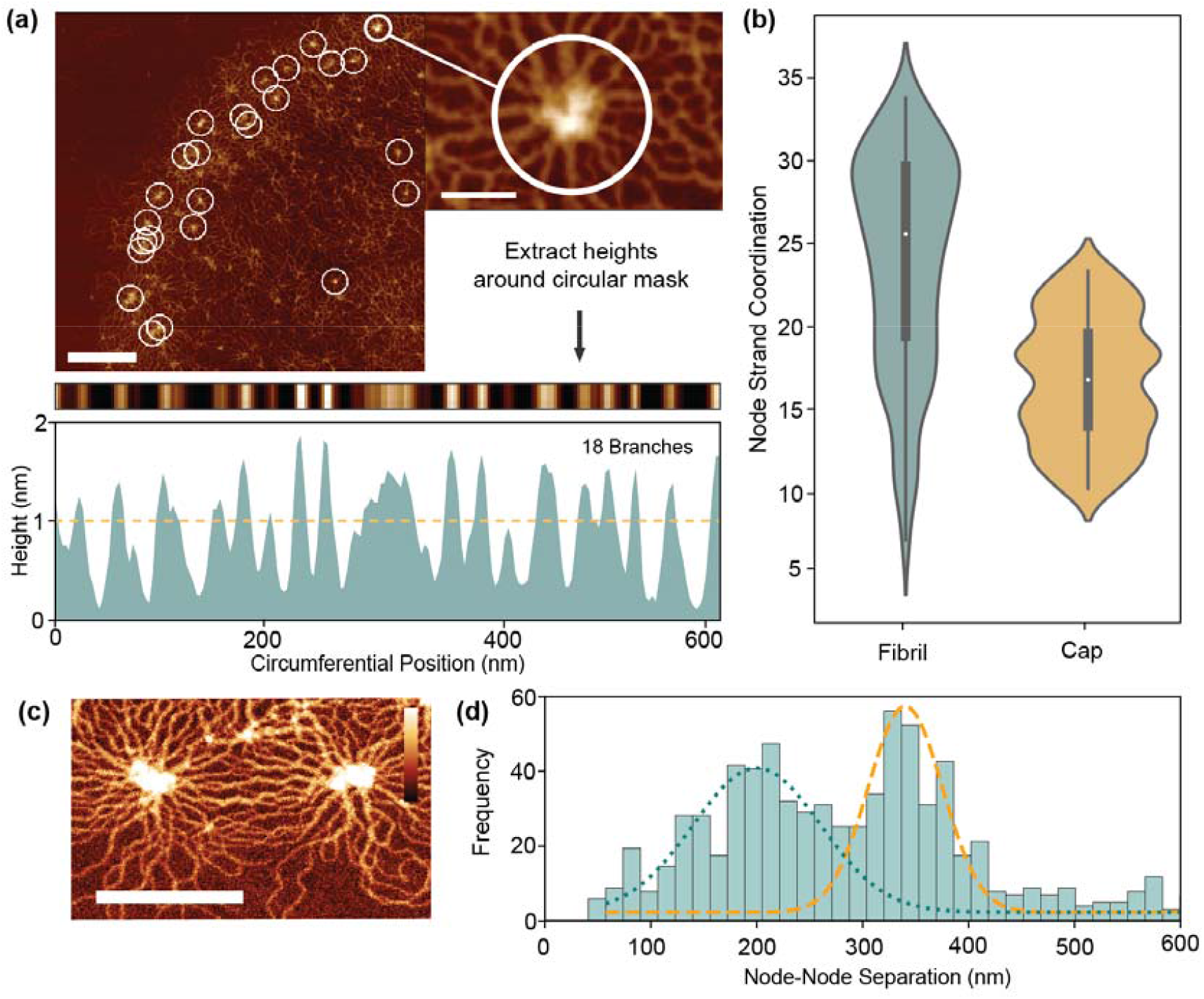
Topological quantifications of network (a) Schematic for tracing of nodes to quantify the connected DNA strands. (b) Distributions for estimated node minicircle coordination between the fibril and cap. (c) AFM image of node sites. Scale bar: 200 nm, height scale: 4nm. (d) Distribution of node-separations over 11 molecules. Bimodal distribution is highlighted with fitting of peaks at 203 nm and 358 nm.

As per Fig. 4b, the mean minicircle coordination of nodes at the fibril is measured to be 26 ± 5, significantly greater than the value for simple valence of 3 which was previously estimated (10,11). This is partly expected as node sites are the highest density points in the kDNA network, so should differ from the mean fibril valence. In the cap, node coordination is generally smaller than in the fibril (17 ± 4), corresponding with the decreased density. These values nonetheless must correspond to the peak coordination with the structure, with catenation at other sites significantly less to maintain the mean of 3. Alternatively, it is feasible that node sites do not represent full catenation between minicircles, but rather a concentration point between partially connected strands. Not all circles must be connected to each other to be held within a tight configuration. As seen in close up images (Fig. 4c), nodes are often an irregular region of varying height and shape. A chain of linking molecules or even physical entanglement, such as knotting, may all contribute to the formation of nodes without full catenation.

To further investigate the nature of nodes, we measured the node separation of all fibril nodes in our dataset. With a mean node separation of 271 ± 14 nm, this broadly corresponds to both the previous findings and theory of He et al. (15), which suggest that the node separation is limited by the half minicircle diameter (425 nm for 2500 bp minicircles). However, a strong bimodal distribution (Hartigans dip test 0.2) is present with peaks at 203 nm and 342 nm. The higher peak likely corresponds to the maximum minicircle separation, with the slight decrease from 425 nm due to the indirect pathways seen in Fig. 4c. However, the lower peak is more unusual and seems to indicate greater complexity within the fibril than previously presumed. This may correlate to the formation of node triples/triangles, where radial stresses cause the formation of a third node between two maximally separated nodes, however this is not observed particularly frequently. It may therefore suggest the presence of more complex phenomena, such as DNA knotting or entanglement between strands.

### Digestion increases molecule curvature and flattens density distribution

The established analytical processes were then applied to kDNA molecules with topological changes induced with two digestion conditions. Using restriction enzymes EcoRI-HF with PstI-HF that digest the minor class of minicircle and maxicircles (22,23), respectively, we remove the minor class minicircles and maxicircles and hence tune the internal topology of the kDNA molecules (named herein as EcoPst). Additionally, the enzyme MluI has been used to partly digest the major class of minicircles and maxicircles (24), which is named herein as M10 due the 10 minute digestion time (Fig. 5a). While the previous in-vitro study of kDNA digestion using these enzymes by Yadav et al. (18) revealed systematic variance in mechanical behaviour (e.g., shape relaxation), the resolution of the experimental technique used limits the determination of structural changes induced by specific class of digestions. Recent work by Ramakrishnan et al. (14) demonstrated the effect of digestion on kDNA molecules with high-resolution AFM, finding that geometric and structural changes occur depending on the class of DNA circle targeted by the enzymes.

**Fig. 5.**
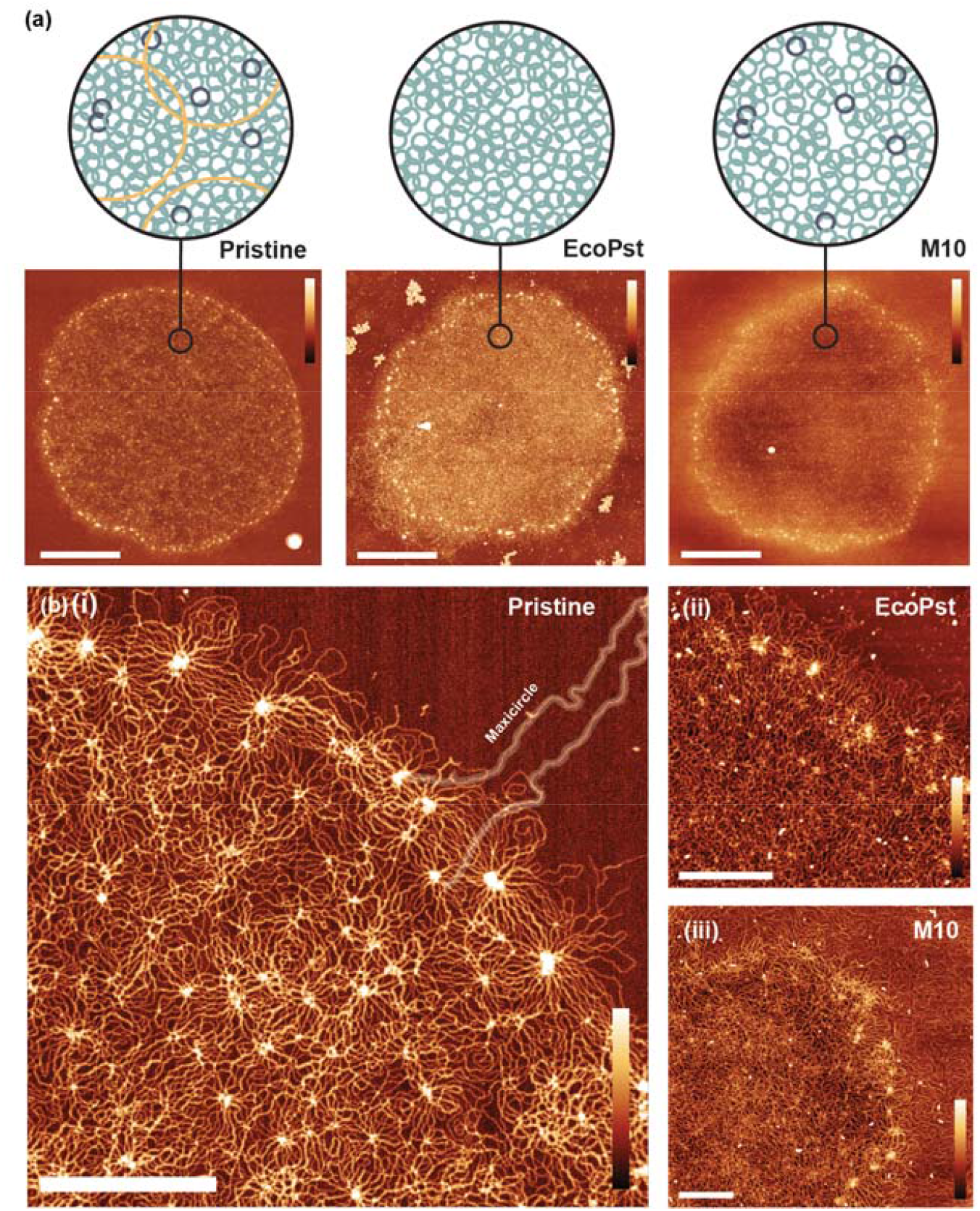
Selective enzymatic digestion of kDNA molecules. (a) Schematic and AFM image for Pristine (no digestion), EcoPst (maxicircle and minor-class minicircles removed), and M10 (maxicircle and ∼14% major minicircles removed). Scale bars: 2 μm. Height Scales: 4 nm (b) Close-up of fibril region. All Height Scales: 3 nm. **i**. Pristine, with maxicircle highlighted. Scale bar: 500 nm **Ii**. EcoPst with less distinct fibril. Scale bar: 200 nm **iii**. M10 with loose fragments visible in the background. Scale bar: 300 nm.

The digestion of the molecules introduced additional complexity to the system, with loose DNA strands and particulates disrupting the quality of imaging. However, as per Fig. 5b, single strand resolution is still achieved in both digestion conditions. As the extent of digestion increases, there is a clear decrease in contrast between the cap and fibril. (Fig. 5). Perimeter, the most consistent geometry of pristine molecules, has a mean 20-30% smaller post-digestion (Fig. 6a). Since the fibril structure remains unbroken after digestion, it suggests that the flattening mechanism is less severe, likely corresponding to a change in the mechanical characteristics of the fibril.

**Fig. 6.**
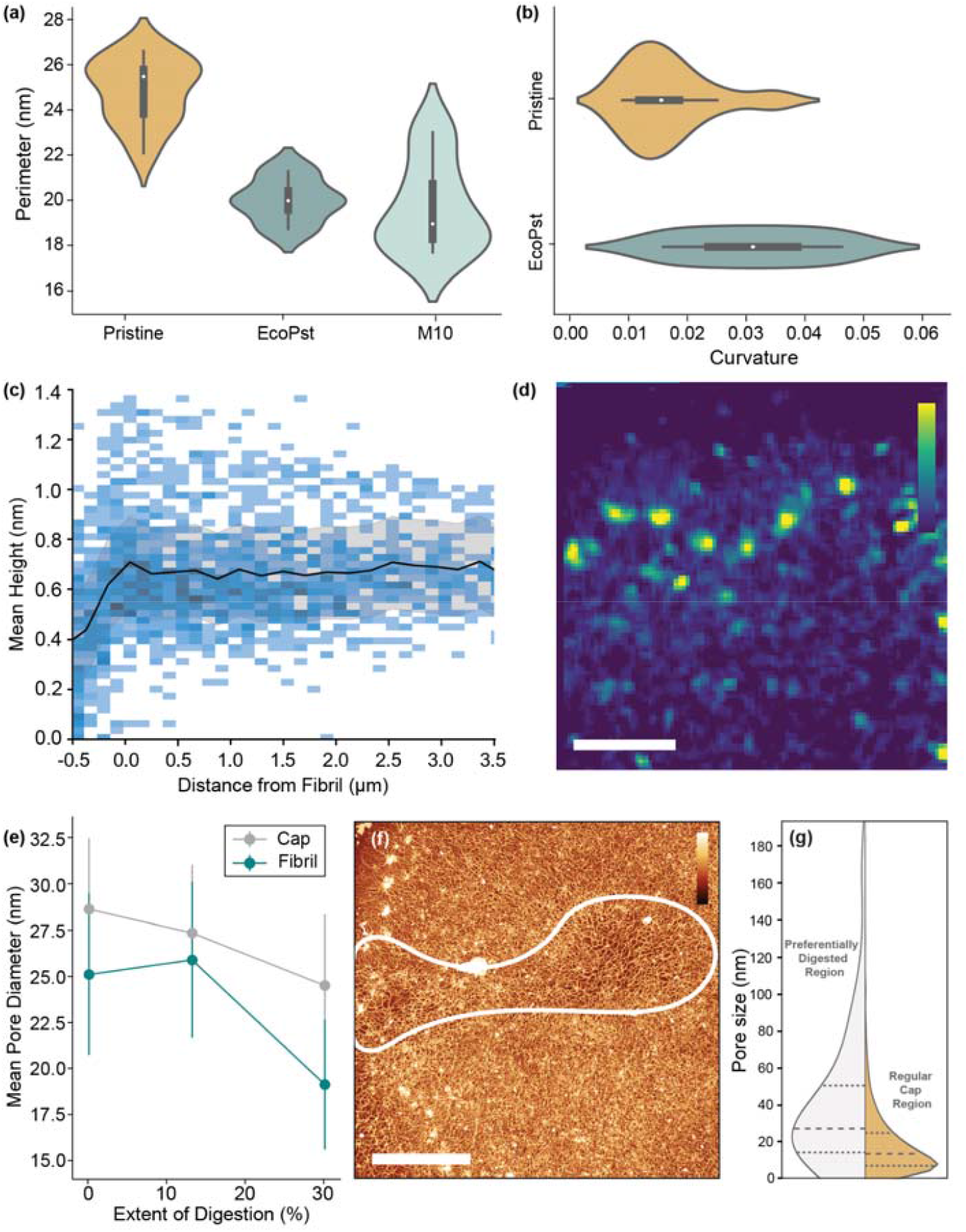
Quantitative analysis of digested kDNA molecules. (a) Violin distributions for perimeter length of molecules with three digestion conditions. (b) Violin plot of curvature for pristine and EcoPst digestion conditions. (c) Mean height distribution and mean value for 6 EcoPst samples. (d) High contrast height images of close-ups of the outer fibril for a single EcoPst molecule. Scale bar: 200 nm. Height scale: 1-4nm. (e) Change in mean pore diameter with extent of digestion for cap and near-fibril positions in EcoPst. (f) Preferential EcoPst digestion of the network, indicated by an increase in local pore size. Scale bar: 1 μm. Height Scale: 3 nm (g) Distribution of pore size through preferentially digested regions.

Subsequently, EcoPst digested molecules show a much increased mean curvature (0.03 ± 0.01) compared to Pristine molecules, indicating that the flattening and stretching process is occurring to a lesser extent (Fig. 6b). As per Fig. 5b, the fibrils of digested molecules have rapid direction changes with less defined sequential arrangement. As the M10 condition displayed a high level of DNA outside the fibril, curvature analysis was not possible for this condition. Furthermore, the fibril was less well-defined, and therefore further analysis could not be performed (Fig. S3).

The density distributions are also affected by digestion, with EcoPst digested molecules having a more consistent mean height throughout (Fig. 6c). Notably, the fibril peak observed at a d = 0 for the pristine molecules (Fig. 2a(ii)), is largely absent from the EcoPst profile, indicating the preferential breakdown of this high-density region. This trend is observed at the smaller length scales (Fig. 6d), with more homogeneous density distributions around the fibril, and none of the trailing behaviour characteristic for pristine molecules. Counterintuitively, the mean height over the molecule is maintained at ∼0.7 nm for the EcoPst digested molecules, in contrast to the decrease in height after the fibril edge to ∼0.4 nm for the pristine molecules. This can likely be prescribed to the reduced size of the molecules upon expansion, increasing the density despite the removal of minicircles, giving a more homogeneous appearance. This trend is also followed by the pore size (Fig. 6e), which generally decreases with the extent of digestion, counter to the expected increase. Upon digestion with EcoPst, molecules are observed to have preferential breakdown in localised areas around the network, both within the cap and the fibril (Fig. 6f). The mean pore size in this preferentially digested region is measured as 37.3 ± 5.7 nm compared to 24.8 ± 4.1 nm in the remainder of the molecule (Fig. 6g). This may correspond to the kinetics of the reaction with enzyme accumulation within a specific region leading to increased digestion rates. However, since this effect is observed over several EcoPst digested molecules, it may correlate to structural heterogeneity. Higher local concentrations of minor-class minicircles could cause less dense areas with greater pore size post-digestion of these components. These results are in agreement with the work by Ramakrishnan et al. who also noted a decrease in size of the molecules upon digestion, as well as a decrease in the circularity (i.e. an increase in curvature) and an increase in the average height after digestion (14).

## Conclusions

We have developed a fully automated pipeline that provides clear insights into the global and local structure of kDNA from raw high-resolution AFM images. This pipeline automates detection of key features such as the cap, fibril and nodes and quantifies the heterogeneous nature of the molecules. Measurements of the network density show the greatest DNA density is located at the fibril node sites around the edge of the molecule, with a steady decrease towards the cap centre. Analysing the global morphology of the molecules, we see that molecules expand significantly on the surface compared to their free state, but that the perimeter remains consistent - indicating that it may regulate the expansion of the molecule. Finally, we utilise our methods to investigate the effect of digestion enzymes on the structure of the kDNA networks. We find a decrease in the molecule size after digestion with an increase in the mean curvature suggest that the flattening and expansion that occurs on adsorption to the surface has been impacted by the digestion. Counterintuitively, we see a decrease in the average pore size, but we propose that this is an effect of the lack of expansion of the molecule. In summary, our high-resolution AFM measurements combined with an automated analysis pipeline provide insight into the structure of the kDNA network and how the individual components may regulate the morphology of the molecule.

## Acknowledgements

This work was supported by a UKRI Future Leaders Fellowship MR/W00738X/1. We wish to acknowledge the Henry Royce Institute for Advanced Materials, funded through EPSRC grants EP/R00661X/1, EP/S019367/1, EP/P02470X/1 and EP/P025285/1 and Robert Moorehead and Xinyue Chen for Dimension FastScan access and support at Royce@Sheffield.

## Data availability

All raw data is publicly available on Figshare (25).

## Code availability

All code is publicly available on Github on the “networks” branch of TopoStats https://github.com/AFM-SPM/TopoStats/tree/SylviaWhittle/networks

## Contributions

B.D., I.Y., E.P.H, P.D. and A.L.B.P., conceived and designed the experiments. B.D. and E.P.H. conducted AFM experiments, B.D. and S.W. wrote software to analyse AFM images and performed analysis. I.Y. prepared samples. D.E.R and T.E.C. supported AFM experiments. T.E.C. supported analysis. T.E.C., P.D. and A.L.B.P. supervised research. B.D. and T.E.C. drafted the manuscript. All authors commented on and revised the manuscript.

## Methods

### kDNA Digestion

kDNA samples, both pristine and digested, were provided in-vitro by Doyle Lab, MIT. The raw kinetoplasts, extracted from *Crithidia fasciculata*, were purchased from TopoGEN Inc. For digestion experiments, different restriction enzymes (EcoRI-HF, PstI-HF and MluI, where HF refers to high fidelity) along with reaction buffer (rCutSmart) were purchased from New England Biolabs, Ipswich, MA. Digestion enzymes (EcoRI-HF, PstI-HF and Mlul) were first mixed with 100μl kDNA samples and left for 2 h, or in the case of Mlul, 10 min at 37°C. The digestion was stopped by heating the enzymes to 80°C for 20 min. Finally, the reaction buffer was replaced by a 0.5X TBE buffer through four-times repeated centrifugation (5000 g, 2 min per run, at 20°C) and by using a 100 kDa cut-off membrane. Samples were shipped at ambient temperature on a next-day service and then stored at 4°C until use.

### AFM Sample Preparation

Synthetic phyllosilicate mica was punched into 1/4” circles and glued directly to a steel sample disk. To prepare the sample, 25 μL of buffer solution (25 mM MgCl_2_, 10 mM TRIS) and 40-60 ng of kDNA were added to a freshly cleaved mica specimen disk. The solution was mixed on the sample disk with 40-50 cycles of the pipette and the DNA was left to immobilise on the mica surface for 5 min. After immobilisation, the sample was rinsed with a heavy stream of MilliQ water for 60 s to remove excess salt and DNA. Finally, compressed air was applied for 5 s to remove the surface water, followed by drying for 10 min in a vacuum chamber (Diener Zepto) at 0.3-0.4 mbar. Vacuum drying under these conditions was selected to cause fast evaporation and prevent the formation of drying defects. The samples were then stored in a desiccator before and between imaging sessions, with samples lasting for up to one month before degradation.

### AFM Imaging

Imaging was performed using the Dimension Icon AFM (Bruker) with PeakForce Tapping mode, with ScanAsyst-Air cantilevers (Bruker). A PeakForce setpoint of 0.01-0.08 V was used which corresponds to peak forces of 440 pN - 3.5 nN. kDNA was first located using a larger scan size of 80-100 μm, and then images were taken over regions between 10 μm and 500 nm. Images were captured at ∼12 nm/px for full molecules and up to 3 nm/px for close-ups.

### Image Processing and Analysis

AFM images were either automatically processed using the TopoStats (19) software (https://github.com/AFM-SPM/TopoStats) or manually with Gwyddion (26). The variation of the code specifically developed to analyse these kDNA images is in the Networks branch of TopoStats. The software first cleans the images by: removing background tilt by fitting a linear polynomial in the x and y directions; performing line by line flattening by setting the median height value of each line to zero; and linearly interpolating through scan line artefacts or scars. To outline the molecule, the kDNA nodes which appear as higher features are used as a guide. A height mask is applied to the image to locate the nodes, with too large or small objects, such as areas of contamination, subsequently excluded. In parallel, a gaussian filter (15 nm) is applied to the original image, followed by a height threshold, which merges all features to produce a general impression of the molecule shape. The outline of this shape is used to filter the detected nodes, with those more than one standard deviation from the outline removed. Together, this provides three levels of conditions for the detection of nodes: their height, size and proximity to the outline of the molecule. This helps to remove the chance of impurity signals disrupting the fitting procedure. The points are sorted based upon their angles around the centroid, then connected point-to-point into a loop which is used to generate geometric parameters such as curvature which are then output to a .csv file. For density measurements, a sliding window of image width/40 (∼250 nm) was used to average the heights over the image, subsampling it. Sub sampled pixels in this image had their distance to the fibril edge measured, allowing for the determination of any correlation between DNA strand density and distance to the edge. Pore size was calculated by applying an Otsu threshold over the image to mask DNA strands. Connected component analysis was used to label each pore, and the square root of each pore’s area was taken as a metric analogous to diameter. Node coordination was performed by identifying nodes via an absolute height threshold; a height profile was taken in a ring around each node as a topological map of the pixel heights around the node. Branches leaving the node were located by using a height threshold, with high regions being identified as branches. The branch count was finally area adjusted and divided by two to give an approximation of minicircle coordination. All code for the algorithms can be found on the Networks branch of TopoStats.

### Statistical Tests

Two-sided Welch’s T-test relation applied to determine the pore size distribution relation between the near-fibril and molecule centre. The t-value was calculated via Eq.1 and the degrees of freedom determined by Eq.2.

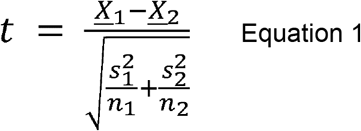

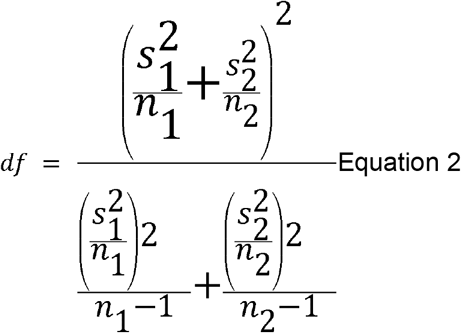

