## Supplementary figures for "Multiscale topological analysis of kinetoplast DNA via high-resolution AFM"

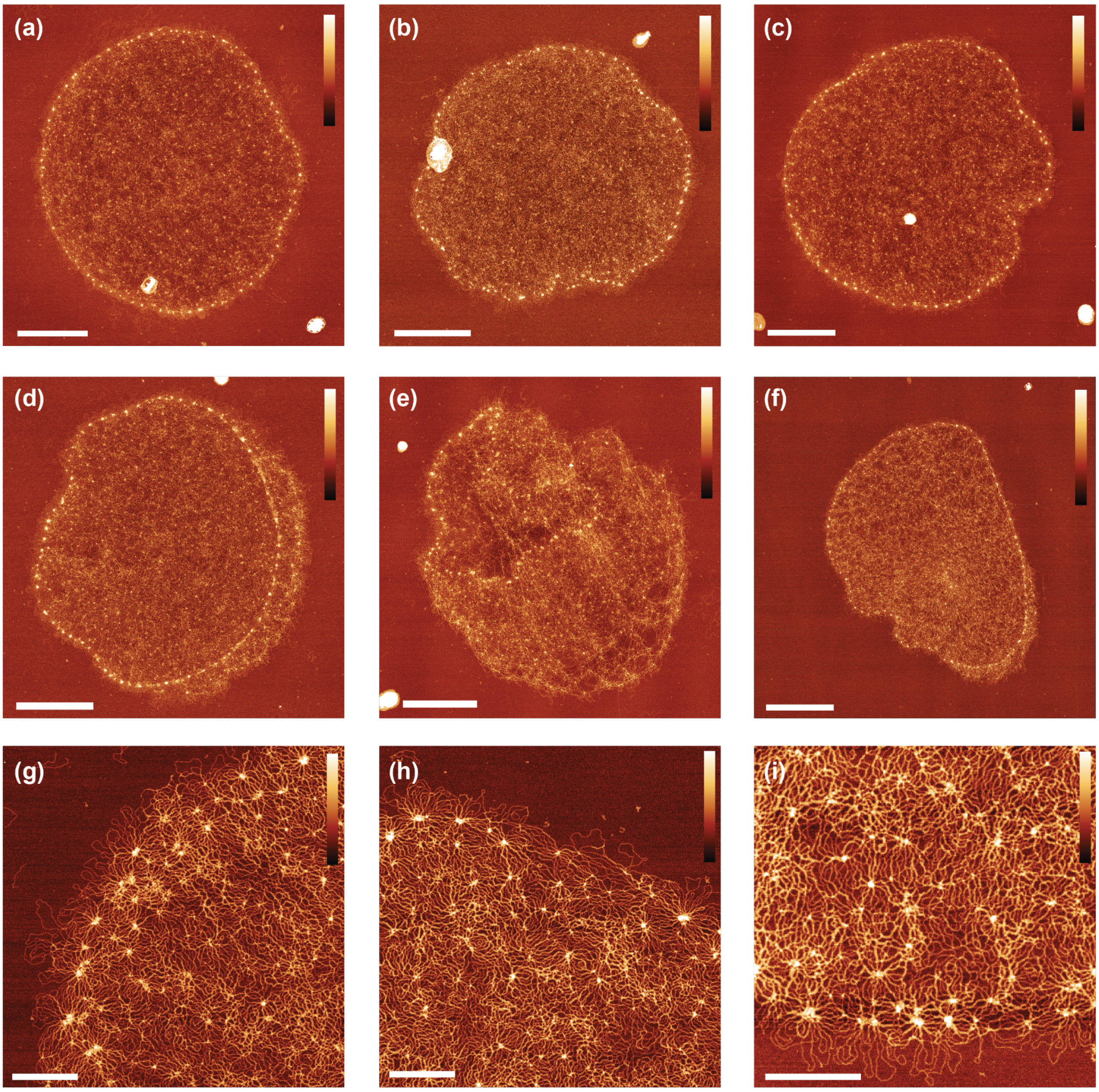


**Fig. S1 -** Compilation of images highlighting the variability observed between molecules of Pristine kDNA. (a-c) kDNA with variable morphology lying flat on the mica surface. Scale bars: 2 µm. Z scale: 4 nm  (d-c) Molecules immobilised partially/fully folded, with the fibril on top of the cap. Scale bars: 2 µm. Z scale: 4 nm (g-i) Close-up images of the fibril region with notable individual character of interior cap. Scale bars: 500 nm. Z scale: 3.5 nm

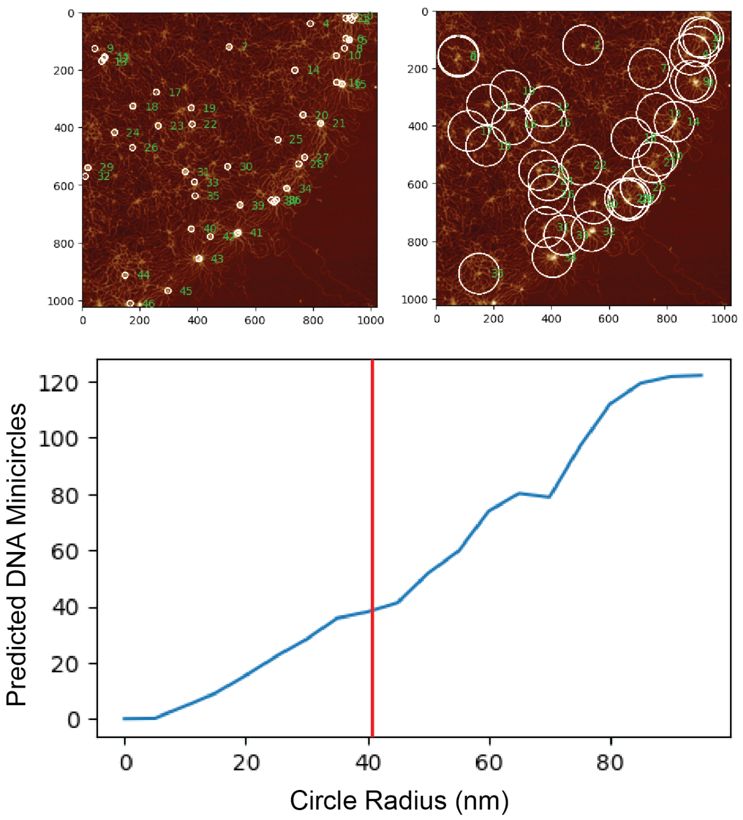


**Fig. S2 -** Node coordination mask size calibration. Masks of 20 nm and 150 nm diameter visualised. Final radius of 40 nm corresponding to plateau after the initial linear region. This value also aligns with the minimum node separation, ensuring the masks intercept as few secondary nodes as possible while achieving maximum resolution.

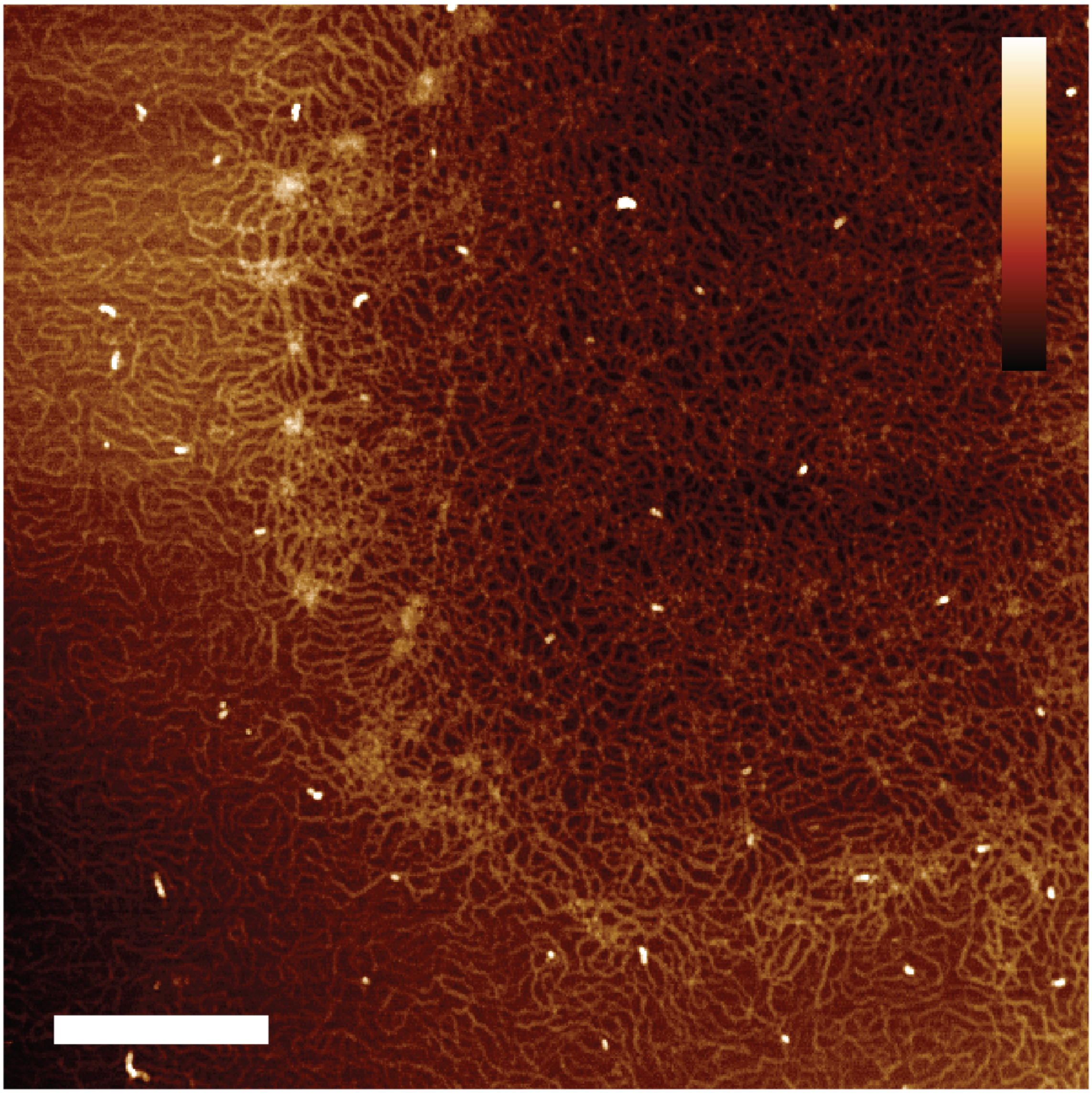

**Fig. S3 -** High levels of background DNA in M10 digested kDNA molecules affect flattening during image pre-processing affecting curvature measurements. Nodes are less visible around the molecule, particularly in the lower region, meaning automated fibril anbaysis could not be performed. Scale bar: 400 nm. Z scale: 4 nm
